# Alignment and mapping methodology influence transcript abundance estimation

**DOI:** 10.1101/657874

**Authors:** Avi Srivastava, Laraib Malik, Hirak Sarkar, Mohsen Zakeri, Fatemeh Almodaresi, Charlotte Soneson, Michael I. Love, Carl Kingsford, Rob Patro

## Abstract

**Background:** The accuracy of transcript quantification using RNA-seq data depends on many factors, such as the choice of alignment or mapping method and the quantification model being adopted. While the choice of quantification model has been shown to be important, considerably less attention has been given to comparing the effect of various read alignment approaches on quantification accuracy.

**Results:** We investigate the influence of mapping and alignment on the accuracy of transcript quantification in both simulated and experimental data, as well as the effect on subsequent differential expression analysis. We observe that, even when the quantification model itself is held fixed, the effect of choosing a different alignment methodology, or aligning reads using different parameters, on quantification estimates can sometimes be large, and can affect downstream differential expression analyses as well. These effects can go unnoticed when assessment is focused too heavily on simulated data, where the alignment task is often simpler than in experimentally-acquired samples. We also introduce a new alignment methodology, called selective alignment, to overcome the shortcomings of lightweight approaches without incurring the computational cost of traditional alignment.

**Conclusion:** We observe that, on experimental datasets, the performance of lightweight mapping and alignment-based approaches varies significantly and highlight some of the underlying factors. We show this variation both in terms of quantification and downstream differential expression analysis. In all comparisons, we also show the improved performance of our proposed selective alignment method and suggest best practices for performing RNA-seq quantification.

## 1 Introduction

Since its introduction in 2008^1–3^, transcriptome profiling via RNA-seq has become a popular and widely-used technique to profile gene- and transcript-level expression, and to identify and assemble novel transcripts. Expression estimation is often done with the goal of subsequently performing differential expression analysis on the gene abundance profiles. In response to improvements in RNA-seq quality and read lengths, as well as significant improvements in the available quantification methods, it has also become increasingly common to perform quantification and differential testing at the transcript level. Recently, very fast computational methods^4–7^ for transcript abundance estimation have been developed which obtain their speed, in part, by forgoing the traditional step of aligning the reads to the reference genome or transcriptome. These methods have gained popularity due to their markedly smaller computational requirements and their simplicity of use compared to more traditional quantification “pipelines” that require alignment of the sequencing reads to the genome or transcriptome, followed by the subsequent processing of the resulting BAM file to obtain quantification estimates.

In various assessments on simulated data^8–10^, these lightweight methods have compared favorably to well-tested but much slower methods for abundance estimation, like RSEM^11^, coupled with alignment methods such as Bowtie2^12^. However, assessments based primarily (or entirely) upon simulated data often fail to capture important aspects of real experiments, and similar performance among methods on such simulated datasets does not necessarily generalize to experimental data. Popular methods for transcript quantification^4–7,11,13–16^ differ in many aspects, ranging from how they handle read mapping and alignment, to the optimization algorithms they employ, to differences in their generative models or which biases they attempt to model and correct. These differences are often obscured when analyzing simulated data, since aspects of experimental data that can lead to substantial divergence in quantification estimates are not always properly recapitulated in simulations.

We focus on the effect of read mapping on the resulting transcript quantification estimates. To compare the effect of different alignment and mapping methods on RNA-seq transcript quantification and related downstream analysis, we have picked tools from three different categories of mapping strategies: (1) unspliced alignment of RNA-seq reads directly to the transcriptome, (2) spliced alignment of RNA-seq reads to the annotated genome (with subsequent projection to the transcriptome), and (3) (unspliced) lightweight mapping (quasi-mapping) of RNA-seq reads directly to the transcriptome. While numerous different lightweight mapping approaches exist^4,5,7,13,17^, and the degree to which they diverge from alignment-based methods can differ, a key feature shared by such approaches is that they do not validate predicted fragment mappings via an alignment score, which precludes them from discerning loci where the best mappings would not admit a reasonable-scoring alignment (i.e. spurious mappings). Furthermore, the focus on speed means that such methods tend not to explore sub-optimal mapping loci, despite the fact that such loci might admit the best alignment scores and therefore be the most likely origin for a fragment. We show that differences in how reads are aligned or mapped can lead to considerable differences in the predicted abundances. Specifically, we find that lightweight mapping approaches, which are generally highly concordant with traditional alignment approaches in simulated data, can lead to quite different abundance estimates from alignment-based methods in experimental data. These differences happen across a large number of samples, but the magnitude of the differences can vary substantially from sample to sample. We also find that these differences appear even when exactly the same optimization procedure is used to infer transcript abundances. Instead, these differences are a result of the different mapping and alignment approaches returning distinct, and sometimes even disjoint, mapping loci for certain reads.

Due to the absence of a ground truth in experimental data, it is difficult to categorically specify which approaches produce more accurate estimates. However, by investigating the divergence we observe among the quantifications produced by different methods, and the differences in read mapping that lead to this divergence in quantifications, we uncover some primary failure cases of different alignment and mapping strategies. This leads us to compare and combine the results of different alignment strategies, and allows us to curate a set of oracle alignments for experimental samples. Comparing various approaches to the oracle provides further evidence for a hypothesis, raised in Sarkar et al.^18^ and Vuong et al.^14^, that lightweight mapping approaches may suffer from spurious mappings leading to a decrease in the resulting quantification accuracy compared to alignment-based approaches. We also demonstrate that, even among alignment-based approaches, non-trivial differences arise between quantifications based upon mapping to the transcriptome (using Bowtie2^12^) and quantifications based upon mapping to the genome and subsequently projecting these alignments into transcriptomic coordinates (using STAR^19^). Both of these alignment-based approaches sometimes disagree with the oracle, but do so for different subsets of fragments and to a varying degree among different samples.

Finally, we introduce an improved mapping algorithm, selective alignment (SA), that is designed to remain fast, while simultaneously eliminating many of the mapping errors made by lightweight approaches. SA is integrated into the Salmon^6^ transcript quantification tool. Our proposed method increases both the sensitivity and specificity of fast read mapping. It relies upon alignment scoring to help differentiate between mapping loci that would otherwise be indistinguishable due to, for example, similar exact matches along the reference. Our approach also determines when even the best mapping for a read exhibits insufficient evidence that the read truly originated from the locus in question, allowing it to avoid spurious mappings.

We also attempt to address one of the failure modes of direct alignment against the transcriptome, compared to spliced alignment to the genome: when a sequenced fragment originates from an unannotated genomic locus bearing sequence-similarity to an annotated transcript, it can be falsely mapped to the annotated transcript since the relevant genomic sequence is not available to the method. We describe a procedure that makes use of MashMap^20^ to identify and extract such sequence-similar *decoy* regions from the genome. The normal Salmon index is then augmented with these decoy sequences, which are handled in a special manner during mapping and alignment scoring, leading to a reduction in such cases of false mappings. We benchmark two variants of this approach; one in which we extract a small collection of decoy sequences via the procedure mentioned above (designated as `SA`), and one in which we align against the transcriptome and whole genome simultaneously, allowing us to detect fragments that better map to a non-transcriptomic target. The latter approach, which essentially treats the whole genome as *decoy* sequence, is denoted as `SAF`. We benchmark these approaches on both simulated data and a broad collection of experimental RNA-seq samples, and demonstrate that they lead to improved concordance with the abundance estimates obtained via quantification following traditional alignment.

## 2 Results

### 2.1 Comparison between various alignment and mapping algorithms

For benchmarking, we used quasi-mapping and SA (with either the similar-sequence decoy regions or the whole genome), both available in the Salmon^6^ program, where quasi-mapping is a representative for lightweight mapping methods and SA is our proposed method that performs sensitive lightweight mapping followed by an efficient alignment-scoring procedure. For unspliced read alignment directly to the transcriptome, we used Bowtie2^12^, which is an accurate and popular tool for unspliced alignment. Similarly, we used STAR^19^ as representative of methods that perform spliced read alignment against the genome. We chose STAR, in particular, since it has the ability to project the aligned reads to transcriptomic coordinates, which allowed us to use a consistent quantification method, and also because it is part of the popular STAR^19^/RSEM^11^ transcript abundance estimation pipeline.

We used Salmon as the main quantification engine in our analyses since it supports quantification from quasi-mapping, SA, and via the output of traditional aligners. To the best of our knowledge, Salmon is the only quantification tool that has support for both lightweight mapping approaches and quantification using traditional alignments. We used Salmon in alignment mode to process the output from Bowtie2 and STAR. In tests on the initial simulated data, we also included RSEM. To remove variability in the quantification methods that is ancillary to our focus on mapping and alignment, we used the `--useEM` flag in Salmon for comparison against the EM-based algorithm of RSEM. Likewise, to eliminate variability due to the target set of transcripts being quantified, we passed the `--keepDuplicates` option to Salmon when indexing for subsequent mapping using quasi-mapping or SA*.

Where mentioned, the “strict” and “RSEM” versions of Bowtie2 and STAR refer to these tools being run with the flags recommended in the RSEM manual^21^, which disallow insertions, deletions and soft-clipping in the resulting alignments. The difference between them is that the “strict” versions are quantified using Salmon and the “RSEM” versions using the RSEM expression calculation method. Throughout the text, we refer to the pipelines by the following shorthand (more details about the methods are given in Section 4.4, and Table S1 and the full command line options provided to each tool are given in Section 4.5):

- Bowtie2 – Alignment with Bowtie2 to the target transcriptome and allowing alignments with indels, followed by quantification using Salmon in alignment mode.
- Bowtie2_strict – Alignment with Bowtie2 to the target transcriptome and disallowing alignments with indels (i.e. using the same parameters as those used by RSEM), followed by quantification using Salmon in alignment mode.
- Bowtie2_RSEM – Alignment with Bowtie2 to the target transcriptome and disallowing alignments with indels, followed by quantification using RSEM.
- STAR – Alignment with STAR to the target genome (aided with the GTF annotation of the transcriptome) and projected to the transcriptome allowing alignments with indels and soft clipping, followed by quantification using Salmon in alignment mode.
- STAR_strict – Alignment with STAR to the target genome (aided with the GTF annotation of the transcriptome) and projected to the transcriptome and disallowing alignments with indels or soft clipping, followed by quantification using Salmon in alignment mode.
- STAR_RSEM – Alignment with STAR to the target genome (aided with the GTF annotation of the transcriptome) and projected to the transcriptome and disallowing alignments with indels or soft clipping, followed by quantification using RSEM.
- quasi – quasi-mapping directly to the target transcriptome, coupled with quantification using Salmon in non-alignment mode.
- SA – Selective alignment directly to the target transcriptome and a set of decoy sequences (high similarity with the transcriptome), coupled with quantification using Salmon in non-alignment mode. (Details in Section 4.2 and Section 4.3.)
- SAF – Selective alignment (full) directly to the target transcriptome and the genome, treated as decoy sequences, coupled with quantification using Salmon in non-alignment mode. (Details in Section 4.3.)

### 2.2 Performance on typical simulations

We used a Polyester^22^-simulated dataset to show the performance of various methods on synthetic data. The distribution of transcript expression for this simulation was learned from an experimental (human) sample (`SRR1033204`, quantified using Bowtie2 with Salmon). We computed the Spearman correlation of quantification estimates from all the pipelines when compared against a known ground truth (in terms of read count). To simulate technical variation, we ran each simulation 10 times using the same input abundance distribution, but varying the random seed used by Polyester.

We observed that, though there are differences in correlation, all pipelines had somewhat similar overall performance on this simulated dataset, with the exception of STAR, which exhibited the lowest correlation. On this data, quasi-mapping performed better than aligning to the genome (and then projecting to the transcriptome) but marginally worse than doing traditional alignment against the transcriptome (both with and without the strict parameters for Bowtie2). We observed that SA performed better than aligning to the transcriptome using Bowtie2, except when quantified using RSEM, though the difference in this scenario is quite small. Finally, SAF performed very similarly to SA in these simulations, though SA, using the smaller decoy set, performed marginally better given that, in reality, all of the reads truly derive from annotated transcripts (with only very minor modifications due to simulated sequencing error).

Overall, the analysis on this synthetic dataset gives an impression that quantifications resulting from the different mapping approaches exhibit similar accuracy, and that all approaches quantify transcript abundances relatively well. While this is true for these simulated data, we show below that this observation does not generalize to experimental data. We posit that this is because, though great advancements have been made in improving the realism of simulated RNA-seq data, these simulations still fail to capture some of the complexities of experimental data. We describe below one particular way in which the realism of the simulated data can be increased by accounting for variations between the sequenced reads and the transcriptome used for quantification.

### 2.3 Performance on simulations from a variant mouse transcriptome

An observation we made from the previous simulation was that disallowing indels using the RSEM parameters (used for strict and RSEM versions) for Bowtie2 and STAR did not adversely affect accuracy compared to using the default parameters of each method. We hypothesized that this is because the reads are simulated exactly from the reference transcriptome that is being used for alignment and quantification, and only sequencing errors (which are taken to consist entirely of substitution errors) are introduced by the simulator. Yet, in experimental data, the sample being quantified likely exhibits variation with respect to the reference against which the reads are aligned. Some of these variants will be single nucleotide polymorphisms (SNPs), while others will be indels and yet others may be larger structural variants. Thus, restricting alignments to disallow indels seems undesirable, unless one is quantifying against a personalized reference that is known to contain the variants present in the sample, which can potentially improve the accuracy of transcript quantification^23,24^.

**Table 1:** Spearman correlation against ground truth for data simulated using Polyester. Note that the counts were based on a real sample from human.

| Method | Truth |
| --- | --- |
| Bowtie2 | $0.940 \pm 0.001$ |
| Bowtie2_strict | $0.941 \pm 0.001$ |
| Bowtie2_RSEM | $0.949 \pm 0.000$ |
| SA | $0.945 \pm 0.000$ |
| SAF | $0.944 \pm 0.000$ |
| quasi | $0.937 \pm 0.000$ |
| STAR | $0.915 \pm 0.001$ |
| STAR_strict | $0.912 \pm 0.001$ |
| STAR_RSEM | $0.919 \pm 0.000$ |

To test the hypothesis that disallowing indels in the alignments will adversely affect quantification accuracy when simulating from a reference transcriptome containing realistic variants compared to the reference, we performed the following experiment. We obtained VCF files from the Sanger Mouse Genomes website^†^ describing the variants present in the PWK mouse strain. Using g2gtools^25^, we generated a copy of the GRCm38.91 transcriptome containing the variants (including indels) present in the PWK strain and simulated reads from this transcriptome. The results presented in the first column of Table 2 show that when reads are aligned against the PWK strain’s reference and indels are disallowed, the quantification estimates are as accurate as those derived from alignments allowing indels, as expected. However, when we aligned the reads back to the original mouse reference transcriptome (version GRCm38.91), we observed that, indeed, Bowtie2 performed better than Bowtie2_jstrict (second column of Table 2), and that, generally, disallowing indels in alignments has a negative effect on quantification accuracy.

**Table 2:** Spearman correlation against ground truth for data simulated using Polyester. Note that the reads were simulated using the reference containing the mouse PWK strain’s variants.

| Method | PWK | GRCm38.91 |
| --- | --- | --- |
| Bowtie2 | $0.939 \pm 0.001$ | $0.934 \pm 0.001$ |
| Bowtie2_strict | $0.939 \pm 0.001$ | $0.923 \pm 0.001$ |
| Bowtie2_RSEM | $0.942 \pm 0.001$ | $0.925 \pm 0.001$ |
| SA | $0.941 \pm 0.001$ | $0.936 \pm 0.001$ |
| SAF | $0.939 \pm 0.001$ | $0.935 \pm 0.001$ |
| quasi | $0.940 \pm 0.001$ | $0.926 \pm 0.000$ |
| STAR | $0.935 \pm 0.001$ | $0.929 \pm 0.001$ |
| STAR_strict | $0.934 \pm 0.000$ | $0.914 \pm 0.001$ |
| STAR_RSEM | $0.937 \pm 0.001$ | $0.916 \pm 0.001$ |

To further analyze the influence of indels on quantification, we aligned the transcript sequences from the PWK strain and the original reference using edlib^26^ and counted the total length of indels in each transcript compared to the unaltered transcript’s original length (we refer to this quantity as the indel ratio). We then sorted the transcripts in descending order by their indel ratios, and evaluated at each cumulative subset, the difference in correlation with the truth between the quantifications using the alignment method and its “strict” variant. We evaluated this quantity increasing the cumulative subsets by 1000 transcripts at each step. We observed that the difference between methods is highest in transcripts that have a larger indel ratio (Figure 1). Hence, the impact of disallowing indels in the alignment can be considerable for reads that originate from transcripts that differ from the reference due to the presence of indels, and this can eventually lead to such transcripts being substantially misquantified.

**Figure 1:**
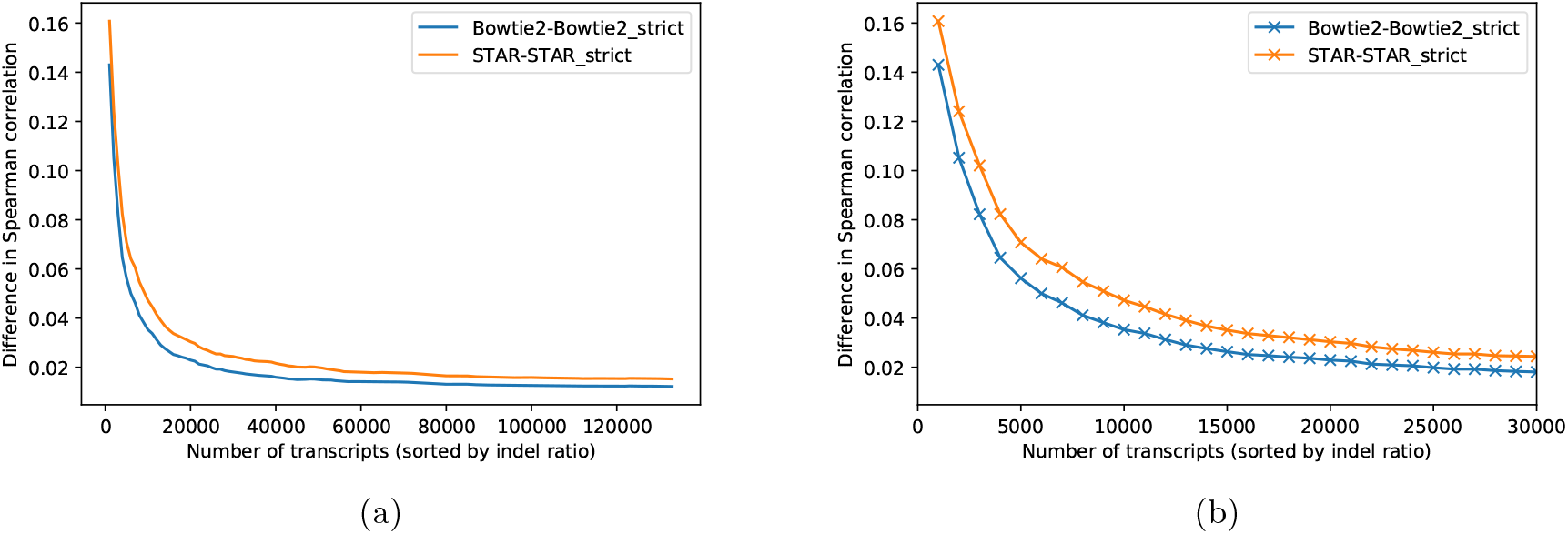
Impact on quantification accuracy when disallowing indels. (a) Difference in correlation with the truth between both alignment methods and their “strict” variants, where indels are disallowed, on all mouse transcripts sorted by their indel ratios. (b) The same plot restricted to the 30,000 transcripts with the largest indel ratio.

Due to both the theoretical concerns and the practical evidence shown here, we proceeded in representing the alignment-based methods by using Bowtie2 and STAR in our comparisons, only in configurations that allow indels to occur in the alignments, and excluded from our analyses the “strict” and “RSEM” versions of the pipelines.

### 2.4 Randomly sampled experiments from NCBI database

It is crucial to evaluate the performance of the various tools on data from real experiments, that can be vastly more complex than even state-of-the-art simulations and can include processes, both known and unknown, that affect the underlying data in complicated ways. To analyze the accuracy of existing tools and study the impact of the artifacts (like the above) on experimental datasets, we randomly selected 200 human RNA-seq experiments from the NCBI database for further investigation. We then filtered the selected samples to include only paired-end libraries having a minimum read length of 75bp. After applying these filters, we were left with a set of 109 samples (69 bulk and 40 full-length single-cell). Before further processing, we applied adapter and (light) quality trimming using TrimGalore^27,28‡^. We also observed that the overall mapping rates across samples tended to be similar between all methods (Figure S1), though Bowtie2 tends to exhibit the highest sensitivity (i.e. aligns the most reads) on average. Subsequently, we quantified all 109 samples using each of the remaining pipelines.

Since no ground truth transcript abundances were available for these 109 experimental datasets, it became more difficult to analyze the accuracy of the different pipelines. However, we explored, manually, some of the cases where differences in mappings and alignments led to divergence of quantification estimates between methods. Between Bowtie2, quasi-mapping, and STAR, the mappings seemed to fall into one of two major categories. In one case, Bowtie2 seemed to be appropriately reporting a more comprehensive set of best-scoring mappings than STAR and quasi-mapping. In the other case, the resulting sequencing fragment seemed to clearly arise from some unannotated region of the genome — either from intronic or intergenic sequence — and it was spuriously assigned by Bowtie2 and quasi-mapping to some set of annotated transcripts (though not always the same set). This led to the following observation: when the fragment truly originates from the annotated transcriptome, Bowtie2 appears the most sensitive and accurate method in aligning the read to the appropriate subset of transcripts, as is also supported by the variant transcriptome simulations from Section 2.3. However, this same sensitivity can sometimes lead Bowtie2 to spuriously align reads to the annotated transcriptome when they are better explained by some other (unannotated) genomic locus. In this latter case, STAR tends to report the correct alignment for the read, and, appropriately refrains from reporting alignments to annotated transcripts. These complications, in which reads are sequenced from underlying fragments that either overlap or are sequence-similar to annotated transcripts, is yet another factor that leads to divergent behavior between different mapping and alignment approaches, but which is not commonly considered in simulation.

These observations led us to combine information from both Bowtie2 and STAR to derive an *oracle* method, that avoids the obvious shortcomings, as listed above, of either of the constituent methods. To derive the oracle alignments in each sample, we used the following approach. First, we aligned the reads for the sample using both Bowtie2 and STAR, and for STAR we retained both the genomic and transcriptomic BAM files (i.e. we considered all of the alignments that STAR was able to produce to the genome, as well as those that it was able to successfully project to the transcriptome). Subsequently, we examined the reads that were aligned to the transcriptome using Bowtie2, and were aligned to the genome using STAR, but which STAR did not project to the transcriptome. For each such read, we examined the best-scoring transcriptomic alignment records produced by Bowtie2 as well as the best-scoring genomic alignment records produced by STAR. We compared the quality of these alignments between the tools by first parsing the extended CIGAR string (the `MD` tag), and assigning a score to each reported alignment. In our scoring scheme, we assigned 1 to every matched base while penalizing soft-clips, SNPs and indels by assigning a score of 0. We reported the score of an alignment as the sum of the number of properly matched bases along the ends of the read. If the transcriptomic alignment of Bowtie2 was of equal or higher quality to the genomic alignment, then we retained the transcriptomic alignment. Otherwise, we marked the fragment’s alignment records for removal. We then processed the original Bowtie2 BAM file for the sample, removing alignments for all fragments that have been marked for removal. The result was a filtered version of the Bowtie2 BAM file in which spurious transcriptomic alignments have been removed. We quantified the sample by providing Salmon with this filtered BAM file, and refer to the resulting quantification estimates as the *oracle* estimates for this sample.

While other complex alignment scenarios may occur, these oracle estimates represent quantification based on the set of alignments that avoid the obvious shortcomings of the different approaches being considered. Specifically, being based on alignment rather than lightweight-mapping, all alignments benefit from the improved sensitivity of Bowtie2’s search procedure and are guaranteed to support a matching of the read to the reference of at least the required quality. Further, since these alignments are derived from Bowtie2, they likely correspond to a correct and comprehensive set of transcripts when the fragment does, in fact, originate from the annotated transcriptome. Finally, in the case where the fragment does not originate from an annotated transcript, and is instead the product of transcription from an unannotated locus, novel splicing, or intron retention, the corresponding alignment records have been removed using information from STAR’s alignment to the genome, so that the fragment is not spuriously allocated to annotated transcripts. Thus, we treated the oracle quantifications as a proxy for the true abundances in the experimental samples.

In terms of Spearman correlation between all methods, we observed the highest average pairwise concordance between the oracle and SAF, in both the bulk and single-cell samples (Figure 2). SAF had the highest rank in order of its correlation with the oracle across the bulk and single-cell samples (histogram of the frequencies of these ranks in Figure 3), and had the closest mapping rate compared to the oracle (Figure S1). Among the alignment-based methods, STAR displayed higher correlation with the oracle — especially in full-length single-cell samples — than did Bowtie2; a reversal of the trend that was observed with respect to the ground truth on the simulated data in Sections 2.2 and 2.3. Further, the overall correlations were lower, and the differences were larger, in the full-length single-cell samples than in the bulk samples. Bowtie2 tended to correlate highly with SA, while STAR tended to correlate highly with SAF, though SA and SAF were, themselves, highly correlated. Also, Bowtie2 and STAR both had a higher correlation with the oracle than they did with each other. This was somewhat expected since the oracle was created by considering the alignments of both Bowtie2 and STAR. However, this also suggested that the manners in which these approaches diverged from the oracle were largely distinct (i.e. they made different types of mistakes in alignment). The quantifications from lightweight mapping exhibited the lowest overall correlation with the oracle. These results were also indicative of the types of divergence between simulated and experimental datasets that we expected to observe. The trend was similar when comparing TPM values after discarding transcripts shorter than 300bp (Tables S2 and S3) and when comparing read counts predicted by each method, instead of TPM, as shown in Figure S2. For the purpose of analyzing the performance of a lightweight mapping algorithm other than quasi-mapping, we also compared the estimates from kallisto^5^ against the other methods on these 109 experimental datasets. It displayed the lowest overall correlations with the alignment-based approaches (Figure S3). This may be due, in part, to the fact that it altered both the quantification and mapping methodology, and because there were no options to control for structural constraints on the reported mappings (i.e. orphaned and dovetailed mappings). Thus, for consistency, we excluded it from the other analyses in the manuscript.

**Figure 2:**
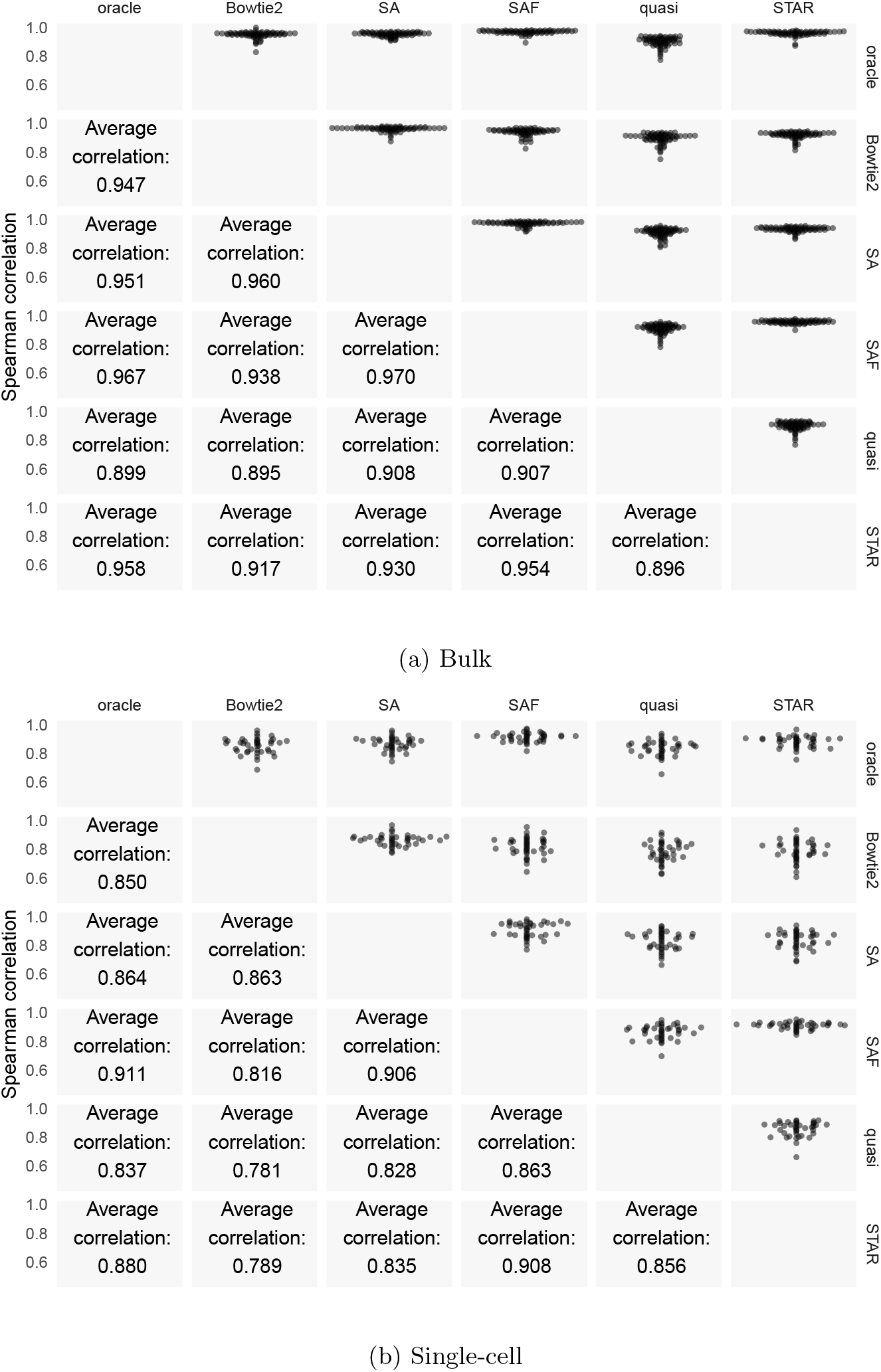
Performance of each method on real bulk and single-cell datasets. The top half of the matrix shows swarm plots of the pairwise correlations of the TPM values predicted by different approaches on the experimental samples. The bottom half shows the average Spearman correlation across the 109 bulk and single-cell samples.

**Figure 3:**
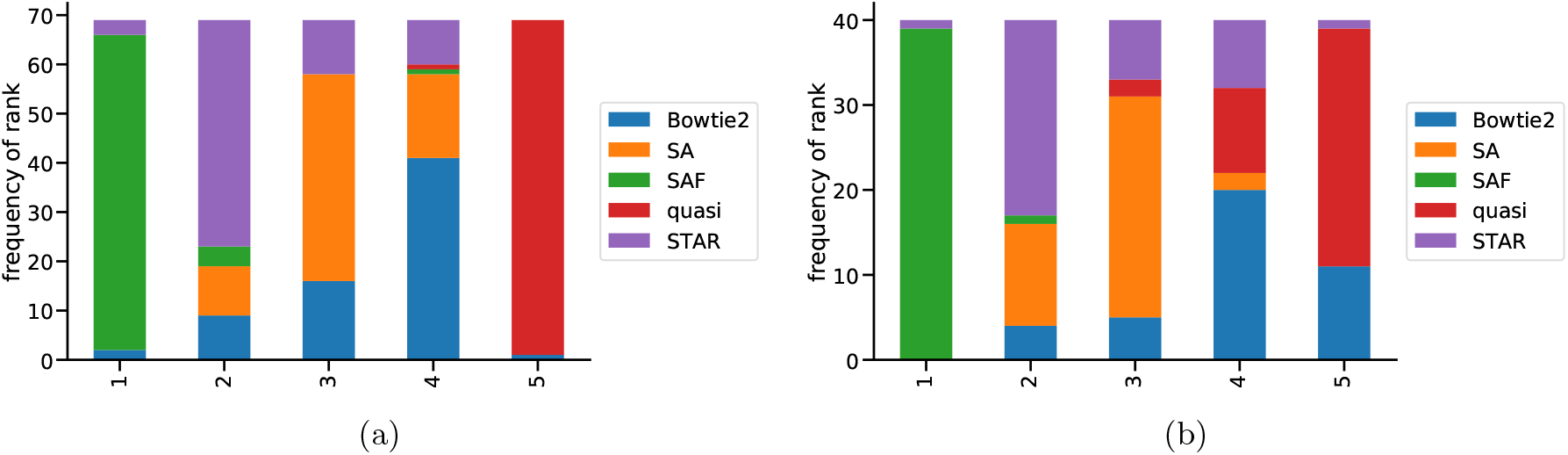
Rank of each method in terms of correlation with oracle. Histogram of the ranks across the 69 bulk samples (a) and the 40 full-length single-cell samples (b), of different methods in terms of the Spearman correlation of the method’s abundance with the oracle. Here, the most correlated method is assigned rank 1, while the least correlated method is assigned rank 5.

Finally, while one would not typically regard any of the average correlations in Figure 2 as poor, it is important to properly frame these differences as representing the aggregate Spearman correlation, across the samples, each quantifying 206, 694 transcripts (wherein most features are properly quantified as having an abundance of 0). A correlation coefficient is a single coarse metric, and as demonstrated in Section 2.3, even substantial quantification differences across thousands of transcripts can nevertheless result in small differences in the summary correlation. To explore some of the transcripts with large differences in quantification across methods, we performed differential transcript expression analysis across methods on the 109 samples, using limma-trend^29^. The counts per million (CPM) for the top 100, 500, and 1000 transcripts is shown in Figure S4. This highlighted the divergence of the methods from each other, in terms of quantification and revealed clusters of transcripts that were differentially expressed under each method. Further, as described in Section 2.6, such differences can lead to considerable changes in which genes are found to be differentially expressed.

### 2.5 Simulation fails to capture complex patterns of real experiments, even when seeded from experimental abundances

In principle, if the specific transcript expression profile was the primary source of quantification difficulty among the different approaches, we should be able to reproduce the types of divergence we observed between different methods in the experimental data (i.e. Figure 2) in simulation by simply creating simulations where the transcript expression profile is seeded with the estimated abundance results obtained from the experimental samples (using e.g. the Bowtie2-derived quantifications). To test this hypothesis, we used Polyester^22^ to simulate 109 synthetic experiments where the expression profiles in each simulated sample were matched to those of the corresponding experimental sample’s transcript abundances generated by the Bowtie2-based pipeline.

We quantified transcript abundance in all of these simulated samples using the same methods we considered in the experimental data. For transcript counts from simulated data, the correlation with the truth, and between methods, is shown in Figure 4. Shown in Figure S5 is the correlation values calculated using the predicted TPMs instead of read counts. Clearly, the correlations among methods was markedly higher in the simulated data than in the experimental data (Figure 2), and multiple methods even showed a correlation ≥ 0.99. The variability between the samples was also considerably lower than what we observed in experimental data. This further suggested that comparison of correlations on simulated data is likely to be only a starting point in assessing different methods, as many salient differences that arise in experimental data disappear when the comparisons are performed on simulated data.

**Figure 4:**
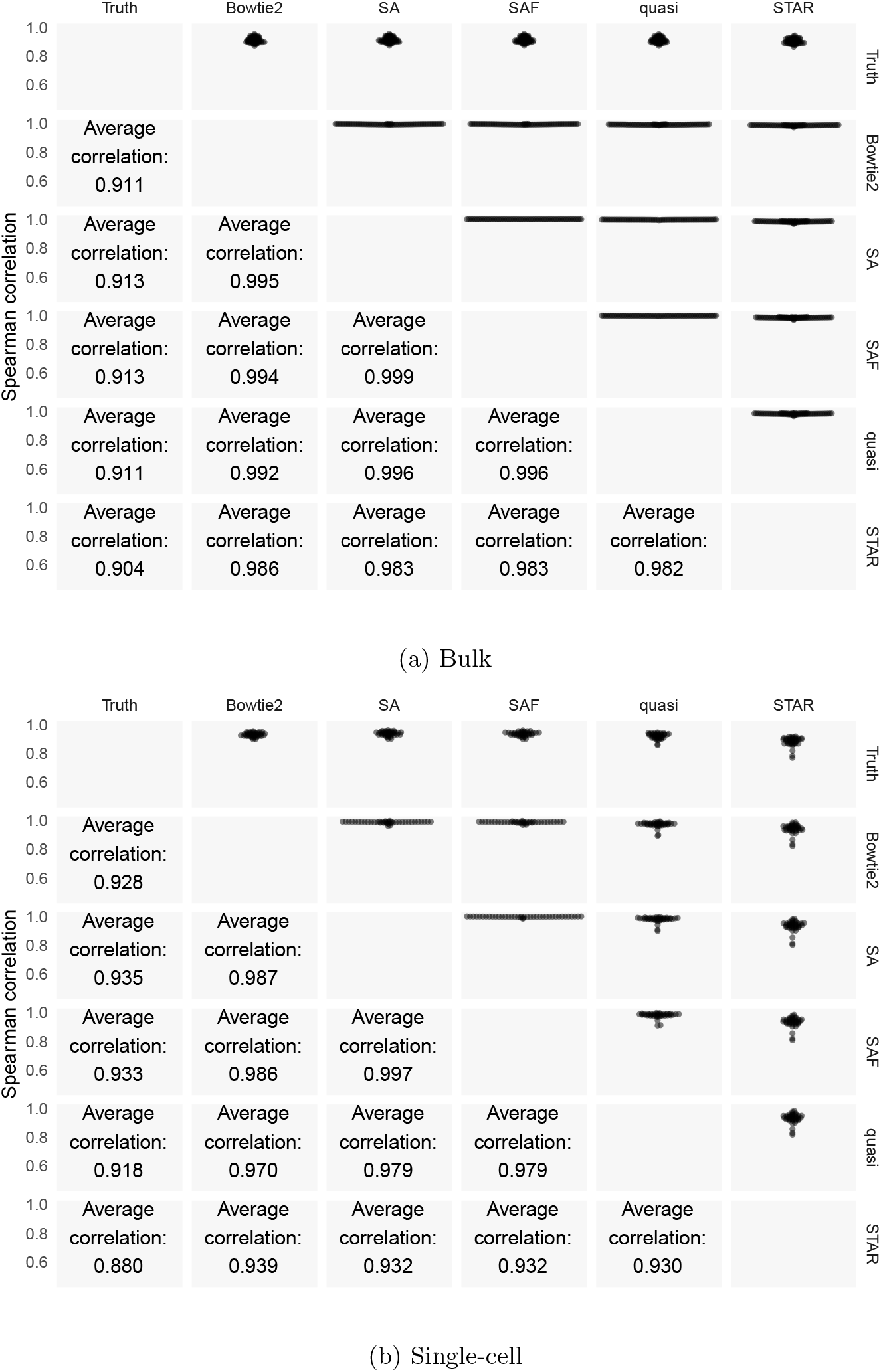
Performance of each method on bulk datasets simulated using Polyester. The top half of the matrix shows swarm plots of pairwise correlations of counts predicted by the different approaches with each other and with the true read counts on the simulated samples generated using the experimentally-derived abundances from Bowtie2 and Salmon. The bottom half shows the average Spearman correlations between the different methods across the 109 simulated datasets.

The variation in correlation across the simulated samples was inconsistent with the hypothesis that the distribution of transcript expressions alone is sufficient to simulate quantification scenarios as complicated as those observed in experimental data. This suggests that there are important aspects, apart from the underlying transcript expression profiles and simulation of read errors, that the simulation failed to capture. Though we do not know all such features, some important biological features, like structural variations (SV), SNP variants and small indel variants, which are sample-dependent rather than reference-dependent, are missing. Furthermore, transcription and sequencing in experimental RNA-seq samples are not limited to the fully-spliced, annotated transcript sequences present in a reference database, even for organisms as well-characterized as human and mouse. Reads derived from unannotated genomic regions that bear resemblance to annotated transcripts, or which only partially overlap the annotated features, can then be spuriously aligned to the annotated features, leading to inaccurate quantification of their abundances. Since such effects can vary from sample-to-sample, they do not affect the estimated expression of the annotated features in a uniform way, and can therefore affect subsequent analyses, such as differential expression testing.

We realize that sample-dependent features are difficult to simulate, and that all of the major features or processes affecting a sample may not even be known, but not including such effects diminishes the realism of the simulated data, and this lack of complexity can be observed in the divergence of the performance of different quantification pipelines compared to how they performed in experimental samples. Identifying other factors present in experimental datasets but lacking in simulations, and determining how to faithfully simulate these factors, seems an important area for future research.

### 2.6 Quantification differences can affect differential gene expression analysis

One of the most common downstream uses of transcript and gene abundance estimates is differential gene expression analysis. Errors in the transcript quantification phase can lead to incorrect detection of differentially expressed genes across conditions. Therefore, quantification is a crucial step for accurate differential gene expression (DGE) estimation and other downstream analyses. To show the influence of quantification on DGE, we performed a case study on two recently published datasets, where sequencing was done for RNA profiling of differences between the healthy and diseased samples. The first dataset is comprised of human ALS motor neurons and studies the effect of the SOD1A4V mutation (2 patient-derived samples) versus (3) isogenic controls (PRJNA236453^30^). The second dataset contained 3 replicates each of uninfected and herpesvirus (HSV-1) infected samples (PRJNA406943^31^). Hence, the design of this study will highlight how misquantifications, possibly arising from incorrect alignments, can impact DGE analysis.

We aligned and quantified reads from all samples using the SA, SAF, quasi-mapping, STAR and Bowtie2 pipelines. The transcript-level counts were summed to the gene level using tximport^32^ and differential expression analysis was performed using DESeq2^33^. Genes were called as differentially expressed between the conditions, for each tool, if they had an adjusted p-value ≤0.01 (i.e. an FDR (false discovery rate) of 0.01 was used). The overlaps of the resulting gene sets were computed. While we had focused on transcript-level analysis in the paper until now, here we looked at differences in gene-level differential expression. This demonstrated that the quantification issues caused by lightweight mapping or misalignment of reads could be of relevance even when one is performing gene-level analyses.

The results, visualized using UpSetR^34^ and presented in Figure 5, show that lightweight mapping tended to miss the largest number of genes discovered as DE by all other approaches (i.e. the second-largest set in both examples, after the consensus set containing genes found by all approaches, was the set of genes found by all approaches except for lightweight mapping). The mean adjusted p-value of these genes under quasi-mapping was 0.034 and 0.069 in the two datasets, showing that lightweight mapping tended to deviate by a large margin from the other methods. Further, lightweight mapping also tended to discover a considerable number of distinct genes that were called as differentially expressed by this approach but not by any of the other approaches (alignment-based or selective-alignment-based), and which may have represented false positives. In each sample, the number of genes with at least one transcript shorter than 300bp constituted less than 10% of the total number of genes called differentially expressed only under the lightweight mapping based quantification, so this effect was unlikely to be driving these differences. In all cases, despite having a large overlap in DE calls with the alignment-based methods, SA produced quantifications that yielded the fewest isolated DE calls. A similar trend was observed when using an FDR of 0.05, as shown in Figure S6(a,b) and when including kallisto as a lightweight mapping method as well, as in Figure S6(c,d). These results suggested that, when the sequenced reads tend to vary more from the reference, as might be the case in many diseased cells, lightweight mapping methods can lead to misquantifications that can eventually lead to false positives and false negatives in downstream differential gene expression studies.

**Figure 5:**
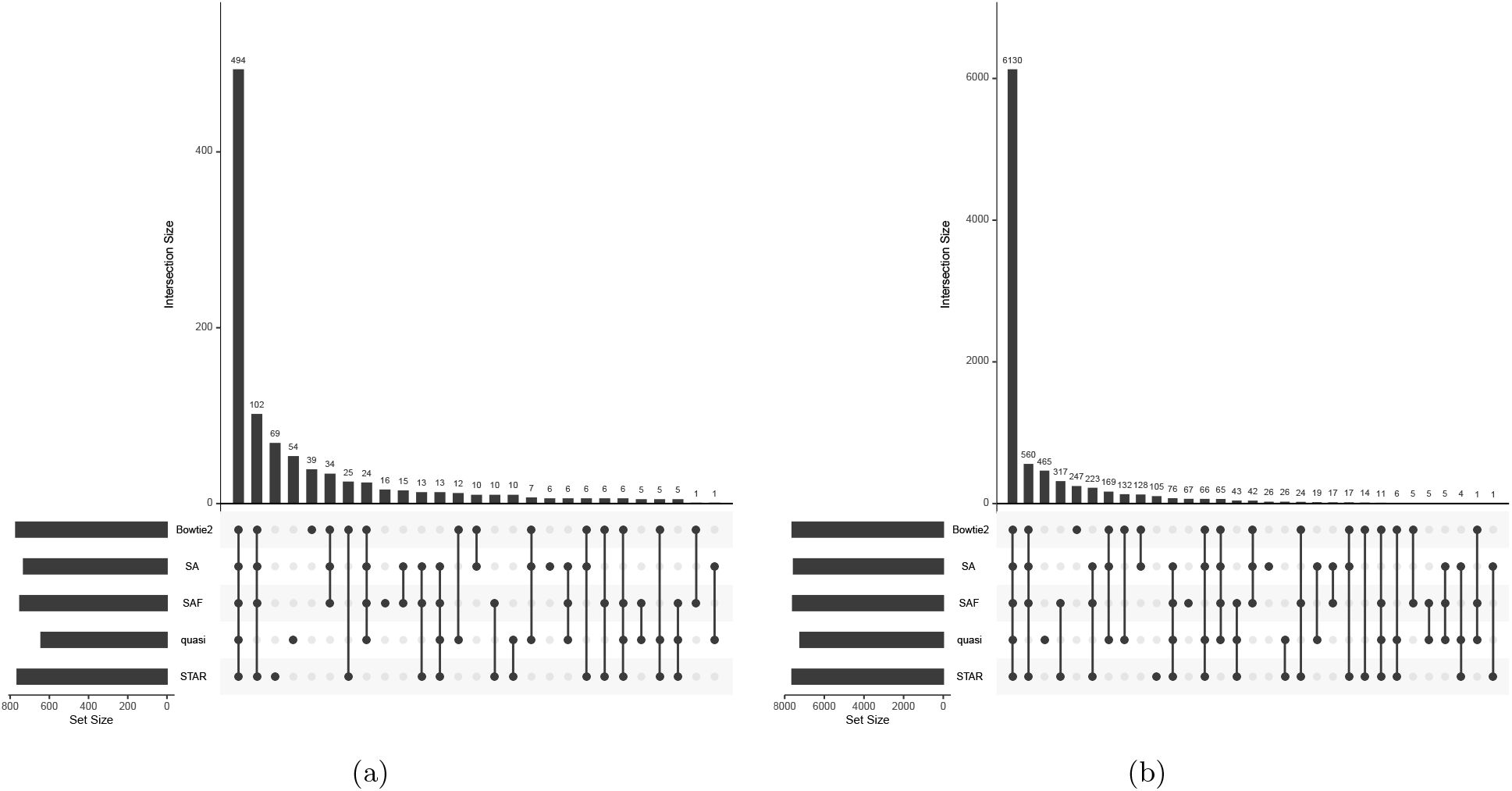
Differentially expressed genes predicted using each method in 2 datasets. Comparison of sets of differentially expressed genes, and their overlaps, computed using each method. The analysis was done on 2 datasets containing multiple replicates of control and infected samples, from human ALS motor neurons (a) and HSV-1 infected cells (b). In each plot, the combination matrix at the bottom shows the intersections between the sets and the bar above it encodes the size of the intersection.

### 2.7 Quantification differences can affect differential transcript expression analysis

Errors in quantification can impact differential expression analysis not just at the gene level, but at the transcript level as well. To show this impact, we selected a dataset that has recently been used to study the impact of the zika virus on the transcriptome in human neural progenitor cells (PRJNA313294^35^). Note that, for consistency, we have used the same design as previous analyses and included both the paired-end and single-end datasets. Hence, there are 2 control and 2 infected samples under each protocol. As with the gene level analysis, the alignment and quantification for all samples was done using the SA, SAF, quasimapping, STAR and Bowtie2 pipelines. Transcript differential expression analysis was done using sleuth^36^ and transcripts were called as differentially expressed at an FDR of 0.01.

The results are presented in Figure 6 and show, as with the gene level analysis, that lightweight mapping tended to be the biggest outlier in terms of missing transcripts called as DE under all other methods. Here, we also observed a considerable set of transcripts (89) discovered only by SAF and STAR, and not by any other approach. For transcripts called as differentially expressed by the other methods but missed by quasi-mapping, the mean adjusted p-value was 0.027. At an FDR of 0.05, this increased to 0.09, highlighting the difference between the quantification estimates from lightweight mapping and alignment-based methods (results visualized in Figure S7(a)). The results presented in Figure S7(b) show the distribution of the DE transcripts if we included kallisto as a mapping and quantification method in this analysis. As before, the lightweight mapping methods, quasi-mapping and kallisto, tended to deviate from the alignment based methods. We also abserved that the number of DE transcripts are the lowest under these methods.

**Figure 6:**
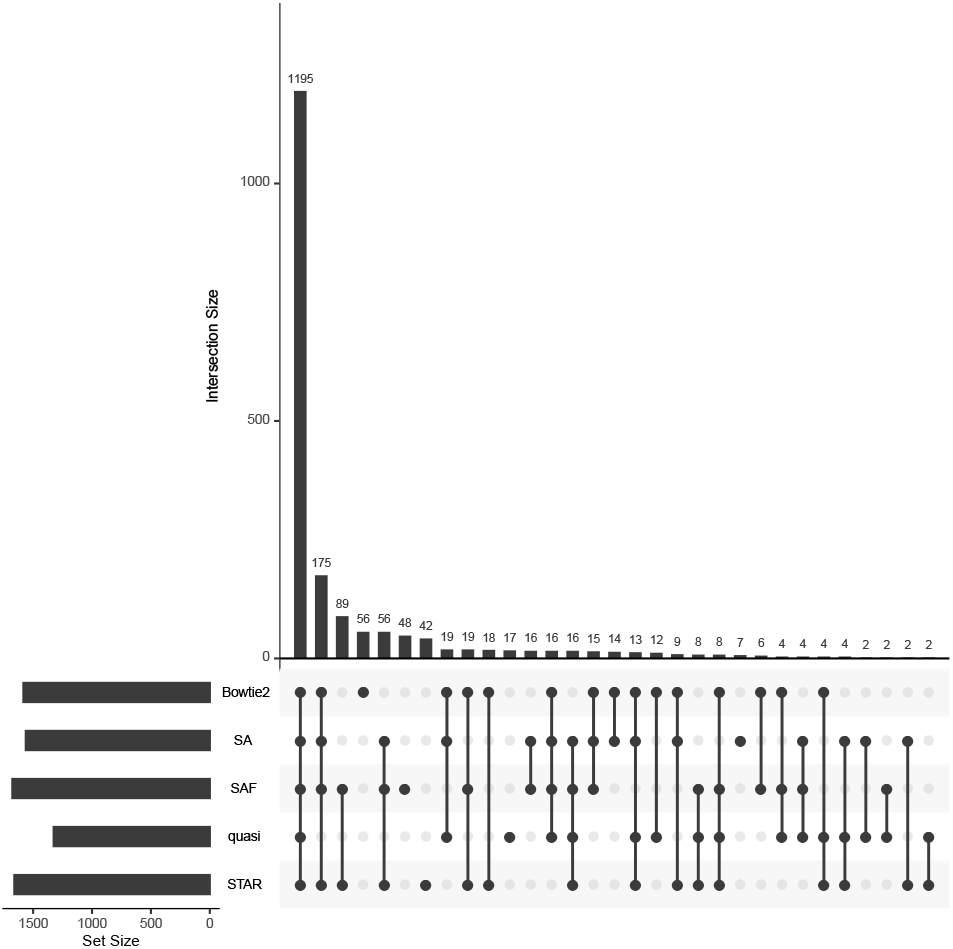
Differentially expressed transcripts predicted using each method. Comparison of sets of differentially expressed transcripts, and their overlaps, computed using each method. The analysis was done on a dataset containing multiple replicates of control and zika infected samples. In each plot, the combination matrix at the bottom shows the intersections between the sets and the bar above it encodes the size of the intersection.

## 3 Discussion & Conclusion

We compared and benchmarked the effects of using different alignment and mapping strategies for RNA-seq quantification, and discussed the caveats implied by different approaches. We observed that methods that perform traditional alignment of the reads against the transcriptome can produce results that are sometimes markedly different from the results produced by lightweight mapping methods. We also observed that performing spliced alignment to the genome and then projecting these alignments to transcriptome can also produce divergent results compared to directly aligning to the transcriptome.

At the same time, we proposed and benchmarked a new hybrid alignment method, SA, which provides an efficient alternative to lightweight mapping that produces results much closer to what is obtained by performing traditional alignment. This approach overcomes the shortcomings of lightweight mapping both in terms of sensitivity and specificity, as it is able to determine appropriate alignments when lightweight approaches return either suboptimal mappings or no mapping, and it is also able to better distinguish the optimal alignment loci among a set of otherwise similar sequences. Some key differences that lead to the improved accuracy of SA are an increase in mapping sensitivity (i.e. more initial mapping loci are explored), a more comprehensive and systematic mechanism for scoring potential mapping loci (making use of the match chaining algorithm of Li^37^), and an actual alignment scoring phase that provides precise information about the quality of each retained mapping, allowing filtering out of spurious mappings that should not be reported. Moreover, the SA approach can take as input a set of decoy sequences, enabling it to avoid some of the spurious transcriptome mappings reported by Bowtie2, when, in reality, the read aligns better to an unannotated genomic locus than to the annotated transcriptome.

The results of benchmarking the different approaches on multiple simulated and experimental datasets lead to a number of conclusions. First, despite the fact that major strides have been made in improving the realism of simulated RNA-seq data, there still remain numerous ways in which simulated data fail to recapitulate the intricacies and challenges of experimental data. One of these is the fact that simulations are almost always carried out on precisely the same transcriptome that is used for quantification, while, in experimental samples, individual variation exists between the sample being assayed and the transcriptome being used for quantification. Another effect not commonly captured in simulation, but prevalent in real data, is the sequencing of reads from unannotated, alternatively-spliced transcripts, from transcripts with retained introns, from otherwise unannotated genomic loci sharing sequence-similarity with annotated transcripts, and from contamination with the sample that may share sequence similarity, to some extent, with the target transcriptome. These effects, along with others that we have not fully characterized in this manuscript, make alignment and quantification in experimental samples much more challenging than in simulated data. Hence, we observed that when quantifying across a broad sample of experimental datasets, the quantification results obtained using different mapping and alignment approaches can demonstrate considerable variation. Together, these results suggest that quantification based purely on lightweight mapping approaches can fail to achieve the accuracy that is obtainable by the same inference algorithms when using traditional alignments, and that these errors in quantification can also affect downstream analyses, even at the gene level (as discussed in Section 2.6). It also suggests that there is practical room for improvement, even in the most accurate existing alignment approaches, at least for the purpose of quantifying the abundance of annotated transcripts.

While it has been previously reported^38^ that pseudoalignment to the transcriptome results in comparable quantification accuracy to alignment to the genome, the analyses performed in this manuscript suggest that alignment to the transcriptome, lightweight mapping to the transcriptome, and alignment to the genome yield quantification results that are sometimes markedly different. There are a few reasons that the analyses carried out in this paper lead to a different conclusion on this question. First, the focus here is much more on experimental as opposed to simulated data. While we found that differences between lightweight mapping and alignment do exist in simulation, the magnitude of their effect on quantification is generally much smaller than is observed in experimental data. Second, while lightweight mapping to the transcriptome and alignment to the genome do yield different quantification results, we also considered traditional alignment to the transcriptome, expanding upon the different common approaches that are taken when aligning reads prior to transcript quantification. Finally, Yi et al.^38^ preprocess both alignments and pseudoalignments into equivalence-class counts (the count of fragments deemed compatible with different subsets of transcripts). Then, from these reduced statistics, abundance estimation is performed. This transformation discards factors that contribute to conditional fragment assignment probabilities like alignment scores (where applicable), fragment lengths, fragment positions, etc. In the analysis presented here, we accounted for such conditional fragment probabilities in the online phase of transcript quantification, and incorporated them (approximately) into the sufficient statistics via the use of range-factorized equivalence classes^39^. Discarding such conditional probabilities could potentially diminish true differences that exist in the underlying mappings that may, depending on the complexity of the quantification model, have an effect on quantification estimates. All of these factors may account for the sometimes considerable differences in quantification accuracy observed downstream of different lightweight mapping and alignment procedures. While we focused on quantification and differential expression, the observations made in this manuscript about the sensitivity and accuracy of different alignment approaches may extend to other downstream analyses as well, such as trans-acting expression quantitative trait locus (eQTL) detection^40^.

Considering only the results on simulated data, one might prefer quantification based on alignment or lightweight mapping of sequencing reads directly to the transcriptome, rather than performing alignment to the genome followed by projection to the transcriptome. One would also observe only small differences between lightweight mapping and alignment to the transcriptome. However, our analyses in experimental data suggested that the increased complexity in real RNA-seq experiments leads to more divergent behavior. In both the bulk and full-length single-cell samples analyzed, SAF yielded the highest overall correlation with the oracle, despite the fact that the oracle is derived from a combination of the Bowtie2 and STAR alignment results. Among the methods based on traditional alignment, alignment to the genome (using STAR, and projecting the resulting alignments to the transcriptome), seemed to display the best concordance, on average, with the quantifications resulting from oracle alignments. SA yielded similar but slightly better accuracy than alignment to the transcriptome using Bowtie2. This is likely, in part, because it is accounting for the sequence-similar decoys that can lead alignment to only the target transcriptome astray. The main benefit of SAF is that it aligns to a reference index that contains both the fully-spliced transcript sequences as well as the entire underlying genome (as potential decoy sequence). This allows SAF to obtain the type of sensitivity that is exhibited by approaches like Bowtie2 and SA when the read truly arises from the annotated transcriptome, but also allows it, like STAR, to avoid spuriously aligning a read to an annotated transcript when it is better explained by some other genomic locus. In the experimental data, both alignment-based approaches and selective-alignment methodologies performed better than quasi-mapping, though the manner in which these methods differ from quasi-mapping, and from each other, were not identical.

When trying to choose an approach, a choice can be made by the user performing the analysis based on any time-accuracy tradeoff they wish to make. In terms of speed, we observed that quasi-mapping is the fastest approach, followed by SA and SAF and then STAR. Bowtie2 was considerably slower than all three of these approaches. However, in terms of accuracy, we found that SAF yielded the best results, followed by alignment to the genome (with subsequent transcriptomic projection) using STAR and SA (using carefully selected decoy sequences). Bowtie2 generally performed similarly to SA, but without the benefit of decoy sequences, seemed to admit more spurious mappings. Finally, lightweight mapping of sequencing reads to the transcriptome showed the lowest overall consistency with quantifications derived from the oracle alignments. The analyses carried out in this manuscript suggest that, with respect to accurate quantification of annotated transcripts, alignment scoring is an important component, but the various pre-existing alignment approaches excelled in different cases. SA takes steps toward addressing the shortcomings of existing alignment based approaches without making large compromises on speed. This is done by indexing parts of the genome that are sequence-similar to the transcriptome or, as in the case of SAF, the entire genome in addition to the annotated transcriptome, hence exhibiting the sensitivity of Bowtie2 in transcriptomic alignment, while avoiding the spurious alignment of reads that do not truly originate from some annotated transcript, like STAR. This approach seemed to provide the highest overall accuracy, at least for the purposes of quantifying an annotated set of transcripts.

## Competing interests

CK and RP are co-founders of Ocean Genomics, Inc.

## Author’s contributions

AS, HS, MZ and RP conceived the idea for the paper. AS, LM, HS, MZ, CS, MIL, RP and CK designed the experiments. AS, LM, HS, CS, MIL and RP carried out the experiments and performed the subsequent analyses. AS, HS, MZ, FA and RP designed and implemented the SA algorithm. All of the authors wrote and approved the manuscript.

## Funding

MIL is supported by R01 HG009937 and P01 CA142538. AS, LM, HS, MZ, FA and RP are supported by R01 HG009937, by National Science Foundation awards CCF-1750472, CNS-1763680 and BIO-1564917 and by grant number 2018-182752 from the Chan Zuckerberg Initiative DAF, an advised fund of Silicon Valley Community Foundation. This work was supported in part by the Gordon and Betty Moore Foundation’s Data-Driven Discovery Initiative [GBMF4554 to CK]; the US National Institutes of Health [R01GM122935, P41GM103712]; and The Shurl and Kay Curci Foundation. The authors thank Stony Brook Research Computing and Cyberinfrastructure, and the Institute for Advanced Computational Science at Stony Brook University for access to the SeaWulf computing system, which was made possible by NSF grant #1531492.

## 4 Materials and Methods

### 4.1 Data and reference

The GENCODE v29 Human reference from https://www.gencodegenes.org/human/release_29.html was used for all experiments involving (simulated or experimental) human reads. The mouse reference genome was obtained from ftp://ftp.ensembl.org/pub/release-91/fasta/mus_musculus/dna/Mus_musculus.GRCm38.dna.toplevel.fa.gz and the GTF was obtained from ftp://ftp.ensembl.org/pub/release-91/gtf/mus_musculus/Mus_musculus.GRCm38.91.gtf.gz. The VCF files for the SNPs and indels were obtained from ftp://ftp-mouse.sanger.ac.uk/REL-1410-SNPs_Indels/mgp.v4.snps.dbSNP.vcf.gz and ftp://ftp-mouse.sanger.ac.uk/REL-1410-SNPs_Indels/mgp.v4.indels.dbSNP.vcf.gz respectively. The list of 109 SRR, scripts to simulate synthetic reads, and the fasta and true abundance files for 10 replicates of simulated data (gencode for human and PWK for mouse) can be found at https://doi.org/10.5281/zenodo.3523437.

### 4.2 Decoy sequences

Alignment against the genome and transcriptome both have their advantages and disadvantages, as discussed earlier. To avoid aligning genomic reads against the transcriptome, without the need to index the complete genome, requires finding regions with high sequence similarity between them. To obtain similar sequences within a reference, we mapped the spliced transcript sequences against a version of the genome where all exon segments were hard-masked (i.e. replaced with `N`). We performed this mapping using MashMap^20^, with a segment size 500 and minimum percent identity of 80%. The sequence similar regions were merged (per-chromosome) using BedTools^41^ and concatenated, giving a decoy sequence for each chromosome. These decoys were then included during the Salmon indexing phase, as described below. A script to obtain these decoy sequences for any reference, given the genome, transcriptome, and annotation is available at: https://github.com/COMBINE-lab/SalmonTools/blob/master/scripts/generateDecoyTranscriptome.sh.

### 4.3 Selective alignment

Selective alignment is based on the pufferfish indexing data structure first described in^42^. Moreover, the index is augmented with the relevant decoy sequence (either restricted decoy sequence as described in Section 4.2 or the entire genome) which is marked during indexing and handled in a special manner during alignment scoring.

The mapping approach works in 5 distinct phases (for paired-end reads). First, exact matches between the read and transcriptome are collected. Second, the set of transcripts to be considered for further processing are extracted. Third, exact match chaining and chain scoring, using the algorithm of Li^37^, is used to determine the relevant putative mapping loci for the read. Fourth (for paired-end reads), the mappings for the first and second read of the pair are matched to determine the mapping loci for the whole fragment. Finally, the computed mapping loci are scored using extension alignment scoring^37,43^ before, after and between the exact matches belonging to the highest-scoring chain for each mapping. In this final step, information about the decoy sequences is used to determine which mappings are considered valid and which are not (details are provided below).

In the first phase of the mapping algorithm, uni-MEMs^44^ are collected between the sequenced fragment (with each read end treated separately) and the index. The uni-MEMs are found via k-mer lookup, and then are extended maximally until the end of a unitig is encountered, the end of the read is encountered, or a mismatch is encountered. If uni-MEM extension terminates because it reaches the end of a unitig or because of a mismatch, search advances in the read and subsequent k-mers are queried in the index to collect other uni-MEMs shared between the read and the index. This process is repeated until the end of the read, and results in a collection of uni-MEMs — matches between the read and the index that can be efficiently decoded into the implied matches between the read and the reference.

In the second phase, uni-MEMs are projected to their corresponded reference loci, and the exact matches are collated by reference and orientation. Let *M* denote the number of matches for the transcript, orientation pair with the maximum number of matches for the current read. An optional (user-determined) filtering policy is applied, whereby any transcript and orientation pair that does not have at least *τM* matches is discarded from further consideration. The value of *τ* is a user-specified fraction, set as 0.65 by default. This optional filtering policy, termed as “filtering before chaining”, is disabled by default, but can be enabled via the command-line option `--hitFilterPolicy BEFORE`. Note that it was not enabled in our experiments and the default was used.

Next, matches along each transcript, orientation pair are sorted and compacted. This compaction is necessary since it is possible to have matches that are directly adjacent on both the read and reference, but which were not extracted as a single exact match during the first phase because the underlying uni-MEM terminated. This compaction phase eliminates such fragmentation due to uni-MEM termination, and reduces the number of exact matches that must be considered by the chaining algorithm. Candidate mapping locations are determined by applying the chaining algorithm of minimap2^37^ to the exact matches for each transcript passing the previous filter. If multiple equally-good positions for a read along a transcript, in terms of their chaining score, are discovered, they are all propagated downstream in the mapping procedure until mappings for paired-end reads are merged. Likewise, if a read is determined to map to a transcript in both the forward and reverse-complement orientation, then all equally-best mapping loci in both orientations are propagated downstream in the mapping procedure. Let *S* be the best chaining score obtained for any mapping of the current read. An optional filter is applied where any mapping with a chaining score less than *τ′S* is discarded from further consideration. By default, this filter, termed as “fitering after chaining”, is enabled by default and *τ′* is set as 0.65.

For paired-end reads, the pairs are merged by determining, for each transcript, the locations of the read ends that respect the expected mapping constraints (e.g. that the leftmost position of the reverse complement read is to the right of the leftmost position of the forward-strand read, and the distances between the reads is less than the maximum allowed insert size). If passed the appropriate flag `--allowDovetails`, then dovetailed^45^ mappings are allowed, but they are prioritized below any non-dovetailed mapping.

All putative mappings are scored using the ksw2^37,43^ library for alignment extension. We note that we compute only the optimal alignment score, and not the details of the alignment itself (i.e. the CIGAR string), which improves the speed of this mapping validation. To avoid redundant computation of the same alignment problem (which is quite prevalent when mapping directly to the transcriptome, as many alignments to alternatively-spliced transcripts will be identical), SA maintains a per-read alignment cache. This alignment cache is a hash table where the key is a hash of the reference transcriptome substring where the read is predicted to align, and the value is the previous alignment score computed for such a substring. Thus, if multiple transcripts would produce identical alignments for the same read, because the read maps to identical regions of these transcripts, SA is able to avoid this redundant work.

Finally, all of the relevant alignments are grouped by their associated alignments scores. Any alignments that fall below the (user-provided) threshold (default of 0.65 of the maximum obtainable alignment score) for a minimum valid alignment score are discarded. During alignment scoring, the score of the best alignment for a given fragment to any decoy sequence as well as to any non-decoy sequence is computed and stored. If the best alignment score to a decoy sequence is strictly greater than the best alignment score to a non-decoy sequence, then all of the fragment’s mappings are considered invalid and the fragment is not considered for quantification. Otherwise, any alignments to decoy sequences are filtered out, and the remaining alignments to valid transcripts are further processed by Salmon using range-factorized equivalence classes^39^, which allows the relevant information about the scores for the different alignments of the read to be appropriately summarized and used for quantification.

### 4.4 Analysis details

#### A note on the difference between SA and SAF

This manuscript introduces the idea of selective alignment over a reference sequence indexed using the pufferfish^42^ data structure, implemented in the Salmon program. The index takes as input a set of decoy sequences, which are not part of the reference transcriptome and are, therefore, not quantified. In our analyses, we considered the performance of the selective alignment algorithm when paired with two different sets of decoys as input. In the first approach, referred to as SA in the manuscript, the index is built on the transcriptome and regions of the genome that have high sequence similarity with the transcriptome. In the second approach, referred to as SAF, the complete genome is included in the pufferfish index as a set of decoys. Hence, the index for SA contains a smaller portion of the genome, whereas the SAF index contains the full genome. Also, note that selective alignment replaces the quasi-mapping algorithm previously used in the Salmon program.

#### A note on on orphan and dovetail mappings

We attempted to normalize for some mapping-related differences between methods that have little to do with the ability of the aligner to appropriately find the correct loci for a read, and instead have to do with constraints placed on what constitutes a valid mapping. Specifically, when projecting to the transcriptome, STAR disallows orphan mappings (cases where one end of a fragment aligns to a transcript but the other end does not). Likewise it is recommended practice in existing alignment-based quantification tools^11,15,16^, when using Bowtie2, to discard discordant and orphaned alignments. Thus, in our analyses, we disallowed orphaned mappings so that, in paired-end datasets, the pair is discarded if only one end of a fragment is mapped, or if the fragment ends only map to distinct transcripts. To be consistent with the default behavior of Bowtie2, the configurations of quasi-mapping and SA were also set to disallow dovetailed mappings (mappings where the first mapped base of the reverse complement strand read is upstream of the first mapped base of the forward strand read). While Bowtie2’s scoring function (when performing global alignment) does not allow insertions or deletions to occur at the beginning or end of the read, we attempted to minimize the effect of this structural constraint on alignments by setting the `--gbar` parameter to 1.

#### A note on genomic alignment, as used in this manuscript

We explored differences that arise between quantification based on alignment of the sequencing reads to the genome and the transcriptome. We considered genomic alignment here to be the process of alignment to the genome — with the benefit of a known annotation — with subsequent projection to the transcriptome. That is, genomic alignment is characterized based on running STAR (with appropriate parameters) to align the reads to the genome, and then making use of the transcriptomically-projected alignments output by STAR via the `--quantMode TranscriptomeSAM` flag (as would be used in e.g. a STAR^19^/RSEM^11^-based quantification pipeline). Such an approach is necessarily concerned only with how well STAR is able to align the sequenced reads to the annotated transcriptome of the organism being assayed, and our assessment is concerned only with the accuracy of quantification of known and annotated isoforms. Importantly, spliced alignment of RNA-seq reads to the genome can be a useful tool in tackling a broader range of problems and in a larger set of cases than can unspliced alignment to a known transcriptome. For example, spliced alignment of sequencing reads to the genome can be done in the absence of an annotation of known isoforms, and can be used to help identify novel exons, isoforms, or transcribed regions of the genome, while unspliced alignment to a pre-specified set of transcripts does not admit this type of analysis. Further, alignment directly to the genome can easily cope with events like intron retention, which are more difficult to account for when using methods that align reads to the transcriptome.

#### A note on the influence of short transcripts on quantification

The human GENCODE v29 reference includes transcripts as short as 8bp, which is much shorter than a single sequencing read or the typical fragment length in most RNA-seq experiments. While RNA-seq might not be the appropriate method to quantify these transcripts, depending on the alignment method, they may have mapped reads and obtain non-zero expression values. In our analyses, we observed that lightweight mapping methods that do not perform end-to-end alignment tend to assign reads to shorter transcripts when there is an exact match. This effect has been explored in some detail by Wu et al.^46^. In such a scenario, it is hard to judge the true origin of the read, and while this may lead to some differences between mapping and alignment-based methods, we showed that the differences in quantification estimates for short transcripts account for only a very small fraction of the overall differences between methods. Since it is difficult to judge how these shorter transcripts, and the reads aligning to them, should be handled, we simply highlighted this issue and refrained from suggesting a particular strategy or attempting to determine which method performed better or worse on transcripts shorter than 300bp.

### 4.5 Tools

We used Salmon v0.15.0 for quasi-mapping and Salmon v1.0 for SA and SAF, Bowtie2 version 2.3.4.3, STAR version 2.6.1b, tximport version 1.12.3, DESeq2 version 1.24.0, kallisto version 0.45.1, edgeR version 3.24.3, limma version 3.38.3, RSEM version 1.2.28, Trim Galore version 0.5.0, bedtools v2.28.0, sleuth version 0.30.0 and MashMap v2.0. All simulated datasets were generated using Polyester version 1.18.0.

For quality trimming the reads we used the following command:

~~~
**trim_galore** -q 20 --phred33 --length 20 --paired <fastq file>
~~~

For indexing, we use the following extra command line arguments, along with the regular indexing and threads parameters:

~~~
**STAR** --genomeFastaFiles <fasta file> --sjdbGTFfile <gtf file> --sjdbOverhang 100
**Bowtie2** default
**salmon**-k 23 --keepDuplicates
**kallisto**-k 23
~~~

For quantification, we use the following extra command line, along with regular index and threads, with each tools we compare against:

~~~
**SA and SAF** --mimicBT2 --useEM
**quasi** --rangeFactorizationBins 4 --discardOrphansQuasi --useEM --noSA
**Bowtie2** --sensitive -k 200 -X 1000 --no-discordant --no-mixed --gbar 1
**Bowtie2_strict** --sensitive --dpad 0 --gbar 99999999 --mp 1,1 --np 1 --score-min L,0,-0.1 --no-mixed --no-discordant -k 200 -I 1 -X 1000
**Bowtie2_RSEM** --sensitive --dpad 0 --gbar 99999999 --mp 1,1 --np 1 --score-min L,0,-0.1 --no-mixed --no-discordant -k 200 -I 1 -X 1000
**STAR** --outFilterType BySJout --alignSJoverhangMin 8 --outFilterMultimapNmax 20 --alignSJDBoverhangMin 1 --outFilterMismatchNmax 999
--outFilterMismatchNoverReadLmax 0.04 --alignIntronMin 20 --alignIntronMax 1000000
--alignMatesGapMax 1000000 --readFilesCommand zcat --outSAMtype BAM Unsorted
--quantMode TranscriptomeSAM --outSAMattributes NH HI AS NM MD
--quantTranscriptomeBan Singleend
**STAR_strict** --outFilterType BySJout --alignSJoverhangMin 8 --outFilterMultimapNmax
20 --alignSJDBoverhangMin 1 --outFilterMismatchNmax 999
--outFilterMismatchNoverReadLmax 0.04 --alignIntronMin 20 --alignIntronMax 1000000
--alignMatesGapMax 1000000 --readFilesCommand zcat --outSAMtype BAM Unsorted
--quantMode TranscriptomeSAM --outSAMattributes NH HI AS NM MD
--quantTranscriptomeBan IndelSoftclipSingleend
**STAR_RSEM** --outFilterType BySJout --alignSJoverhangMin 8 --outFilterMultimapNmax
20 --alignSJDBoverhangMin 1 --outFilterMismatchNmax 999
--outFilterMismatchNoverReadLmax 0.04 --alignIntronMin 20 --alignIntronMax 1000000
--alignMatesGapMax 1000000 --readFilesCommand zcat --outSAMtype BAM Unsorted
--quantMode TranscriptomeSAM --outSAMattributes NH HI AS NM MD
--quantTranscriptomeBan IndelSoftclipSingleend
**RSEM** default
**kallisto** default or --rf-stranded as appropriate
~~~

## Footnotes

* Though we performed indexing here with `--keepDuplicates` and quantification with `--useEM`, this is done only to eliminate controllable sources of variability between methods so as to isolate, as much as possible, the effect of differences in mapping. We generally recommend that duplicate transcripts are discarded during indexing, and that the offline phase of quantification is performed using the variational Bayesian EM.

† ftp://ftp-mouse.sanger.ac.uk/REL-1410-SNPs_Indels/

‡ The effect of trimming on the overall results was relatively minimal (result not shown).

## Notes

https://doi.org/10.5281/zenodo.3523437

