## Supplementary File for "Alignment and mapping methodology influence transcript abundance estimation"

### Supplementary Material for "Alignment and mapping methodology influence transcript abundance estimation"

| | Genome indexed | Alignment scoring | Indels allowed | End-to-end alignment | Quantification method |
|---|---|---|---|---|---|
| Bowtie2 | ✗ | ✓ | ✓ | ✓ | Salmon |
| Bowtie2_strict | ✗ | ✓ | ✗ | ✓ | Salmon |
| Bowtie2_RSEM | ✗ | ✓ | ✗ | ✓ | RSEM |
| STAR | ✓ | ✓ | ✓ | ✓ | Salmon |
| STAR_strict | ✓ | ✓ | ✗ | ✓ | Salmon |
| STAR_RSEM | ✓ | ✓ | ✗ | ✓ | RSEM |
| quasi | ✗ | ✗ | ✓ | ✗ | Salmon |
| SA | ✗* | ✓** | ✓ | ✗ | Salmon |
| SAF | ✓ | ✓** | ✓ | ✗ | Salmon |

Table S1: Various factors altered under each pipeline. *Here, under SA, only regions of the genome that are sequence similar to the transcriptome are indexed, but not the whole genome. Refer to Section 4.2 for further details on how the sequences are obtained. **While SA and SAF produce alignment scores, they do not perform backtracing or reconstruct the edit operations that were used to obtain the optimal alignment score.

| | Oracle | Bowtie2 | SAF | SA | quasi | STAR |
|---|---|---|---|---|---|---|
| Oracle | 1.000/0.000 | 0.858/0.054 | 0.920/0.029 | 0.873/0.046 | 0.843/0.053 | 0.889/0.039 |
| Bowtie2 | – | 1.000/0.000 | 0.817/0.063 | 0.863/0.039 | 0.783/0.068 | 0.791/0.069 |
| SAF | – | – | 1.000/0.000 | 0.907/0.052 | 0.865/0.050 | 0.909/0.023 |
| SA | – | – | – | 1.000/0.000 | 0.829/0.063 | 0.837/0.059 |
| quasi | – | – | – | – | 1.000/0.000 | 0.858/0.052 |
| STAR | – | – | – | – | – | 1.000/0.000 |

Table S2: Mean/standard deviation of Spearman correlation between all methods on 40 single-cell experimental datasets after removing short transcripts with length $< 300$.

| | Oracle | Bowtie2 | SAF | SA | quasi | STAR |
|---|---|---|---|---|---|---|
| Oracle | 1.000/0.000 | 0.949/0.023 | 0.971/0.008 | 0.954/0.014 | 0.907/0.031 | 0.962/0.012 |
| Bowtie2 | – | 1.000/0.000 | 0.940/0.026 | 0.961/0.017 | 0.901/0.034 | 0.920/0.025 |
| SAF | – | – | 1.000/0.000 | 0.972/0.015 | 0.913/0.031 | 0.955/0.011 |
| SA | – | – | – | 1.000/0.000 | 0.914/0.029 | 0.932/0.017 |
| quasi | – | – | – | – | 1.000/0.000 | 0.903/0.029 |
| STAR | – | – | – | – | – | 1.000/0.000 |

Table S3: Mean/standard deviation of Spearman correlation between all methods on 69 bulk experimental datasets after removing short transcripts with length $< 300$.

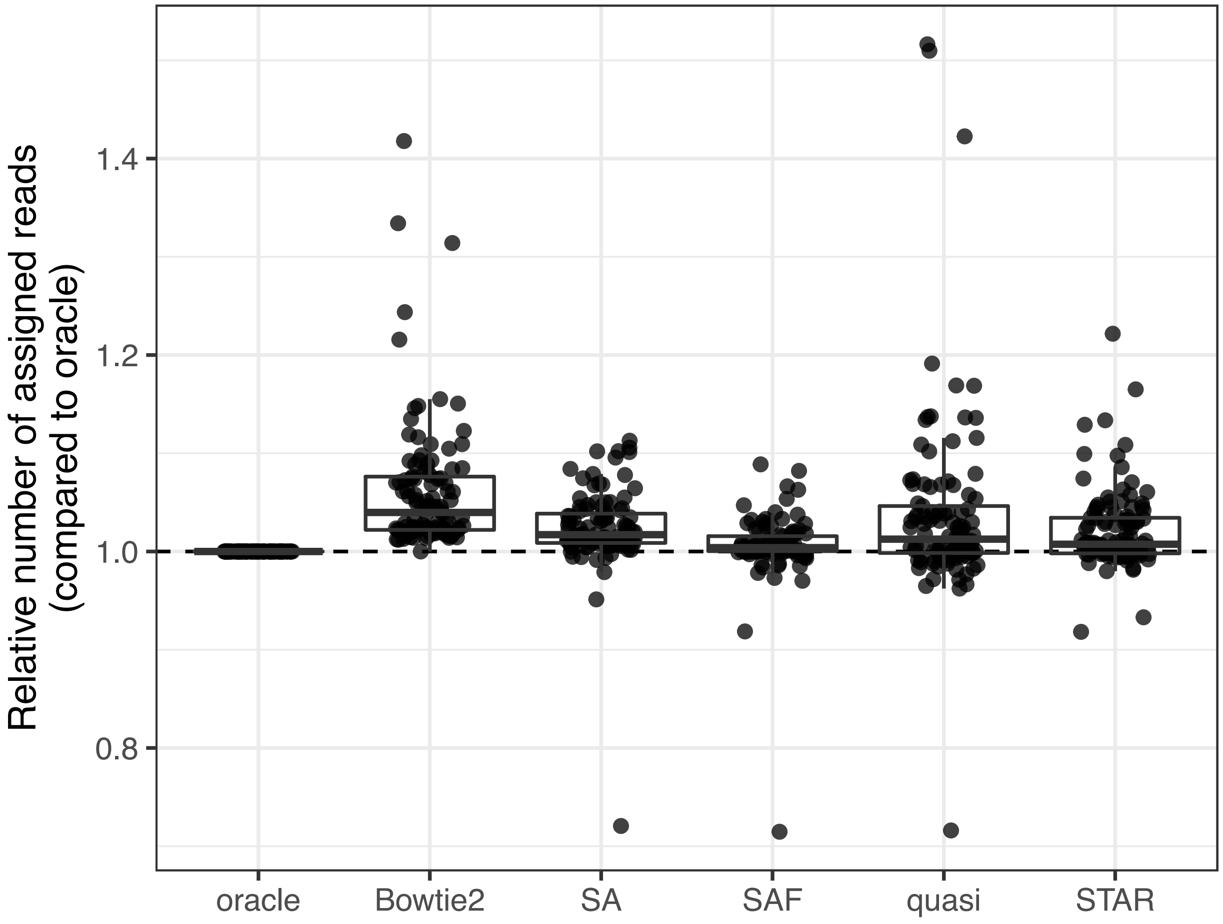


Figure S1: Mapping rates of different methods, relative to oracle, for 109 experimental samples.

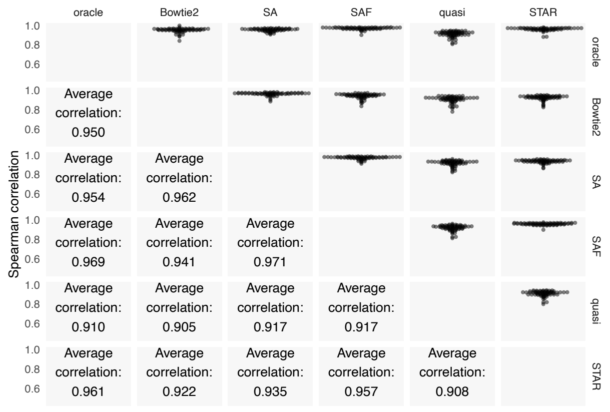


(a) Bulk

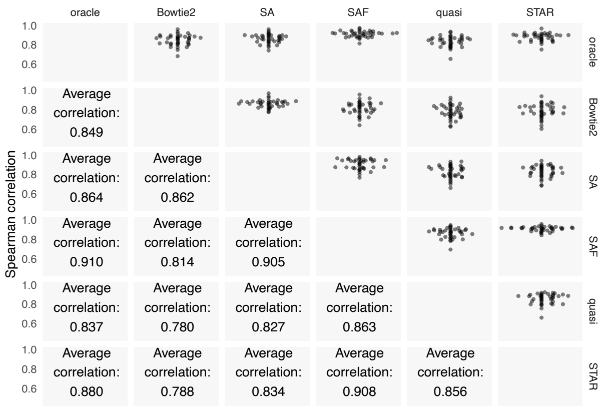


(b) Single-cell

Figure S2: The upper triangle of the matrix shows swarm plots of pairwise correlations of read counts predicted by the different approaches on the experimental samples. The bottom half shows the average Spearman correlations between methods across the 109 bulk and single-cell samples.

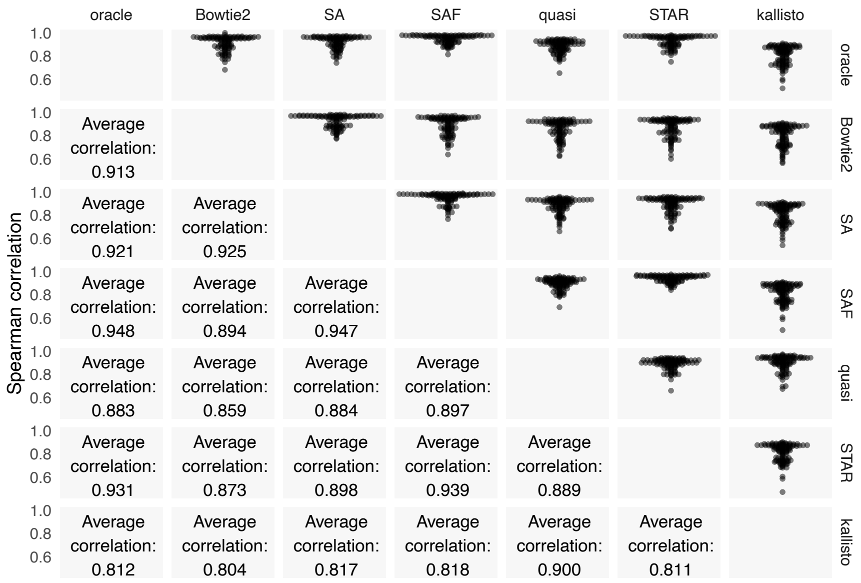


(a)

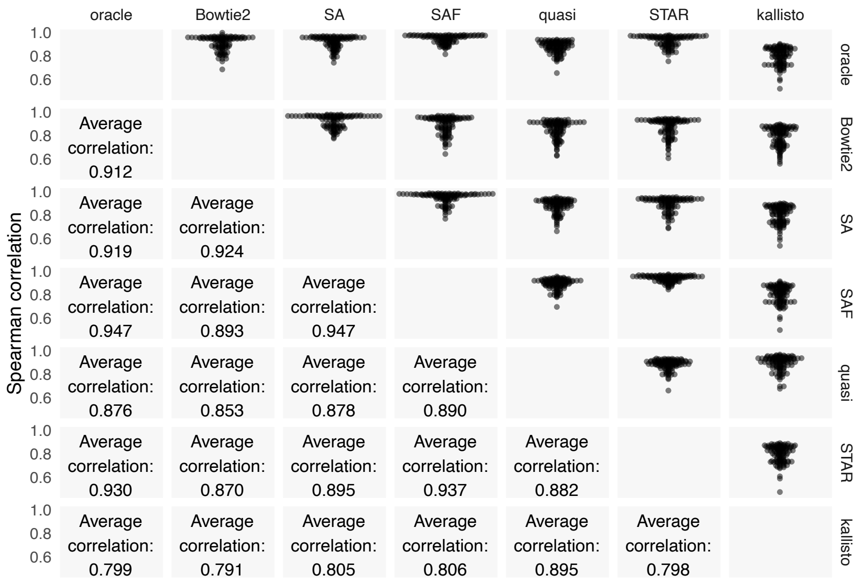


(b)

Figure S3: The top half of each matrix shows swarm plots of the pairwise correlations using count (a) and TPM (b) values predicted by the different approaches on the experimental samples. The bottom half shows the average Spearman correlations between methods across the 109 samples. Note that the quantification method for each pipeline is the same, except kallisto, where both the mapping and quantification algorithms are different. Hence, while other methods disallow orphaned reads and dovetailed mappings, the kallisto output will include them, which may explain, in part, the increased divergence from the alignment-based methods.

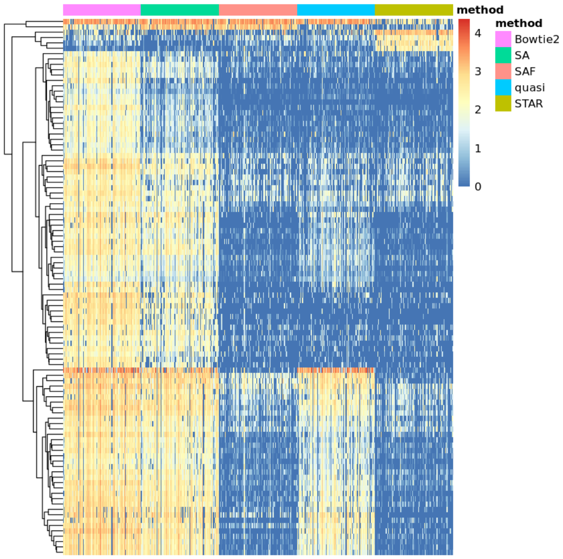


(a)

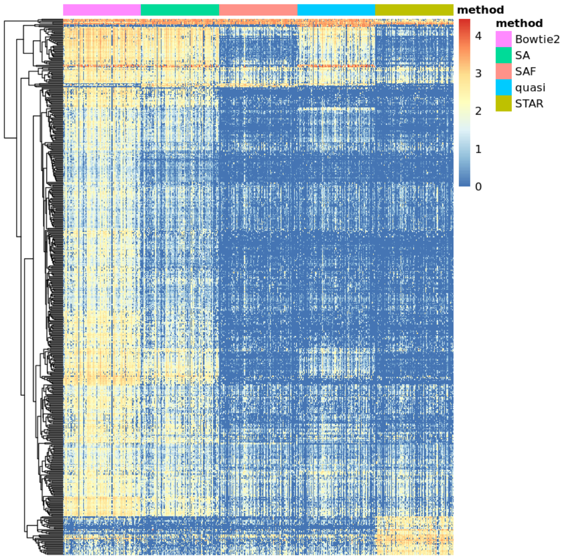


(b)

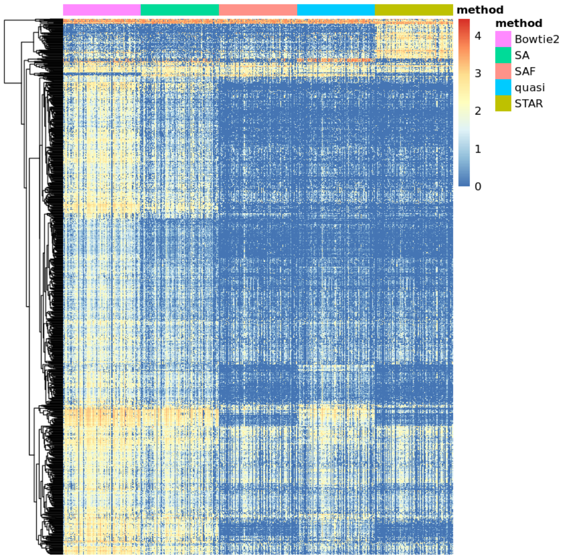


(c)

Figure S4: The $\log_2$(CPM) for 109 samples grouped by method for the top 100 (a), 500 (b), and 1000 (c) differential transcripts. Limma-trend was used with `scaledTPM` counts (generating counts from per-sample TPMs by scaling to the library size) via tximport[32], with a `prior.count` of 3, and using a design of ∼`sample + method`. An F-statistic was generated by specifying coefficients representing differences among the methods and the top transcripts chosen using the F-test p-value.

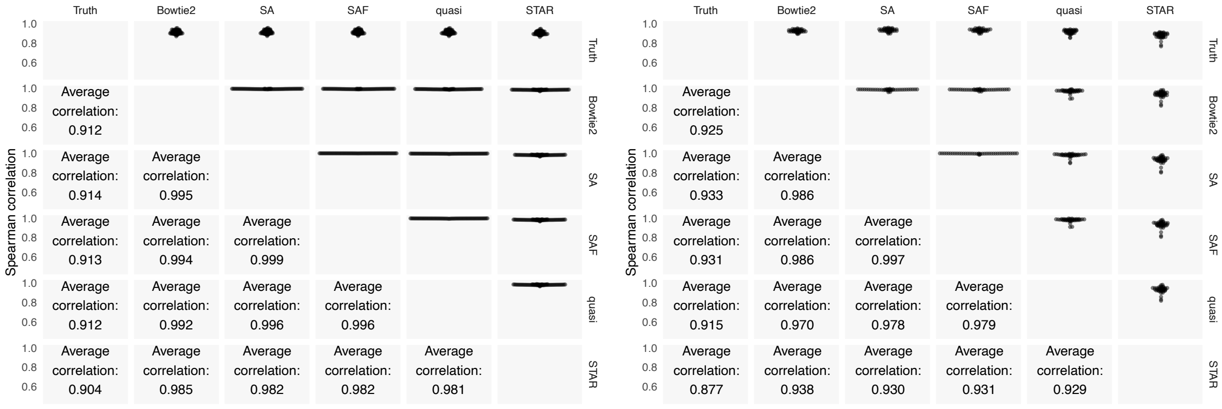


(a) Bulk (b) Single-cell

Figure S5: The top half of the matrix shows swarm plots of the pairwise correlations of TPM values predicted by the different approaches with each other and with the ground truth abundances on the simulated samples. The bottom half shows the average Spearman correlations between the different approaches across the 109 samples. The expected effective length of each transcript was computed according to the true fragment length distribution. Given the true fragment counts and expected effective lengths, the TPM is computed as in Li and Dewey[11].

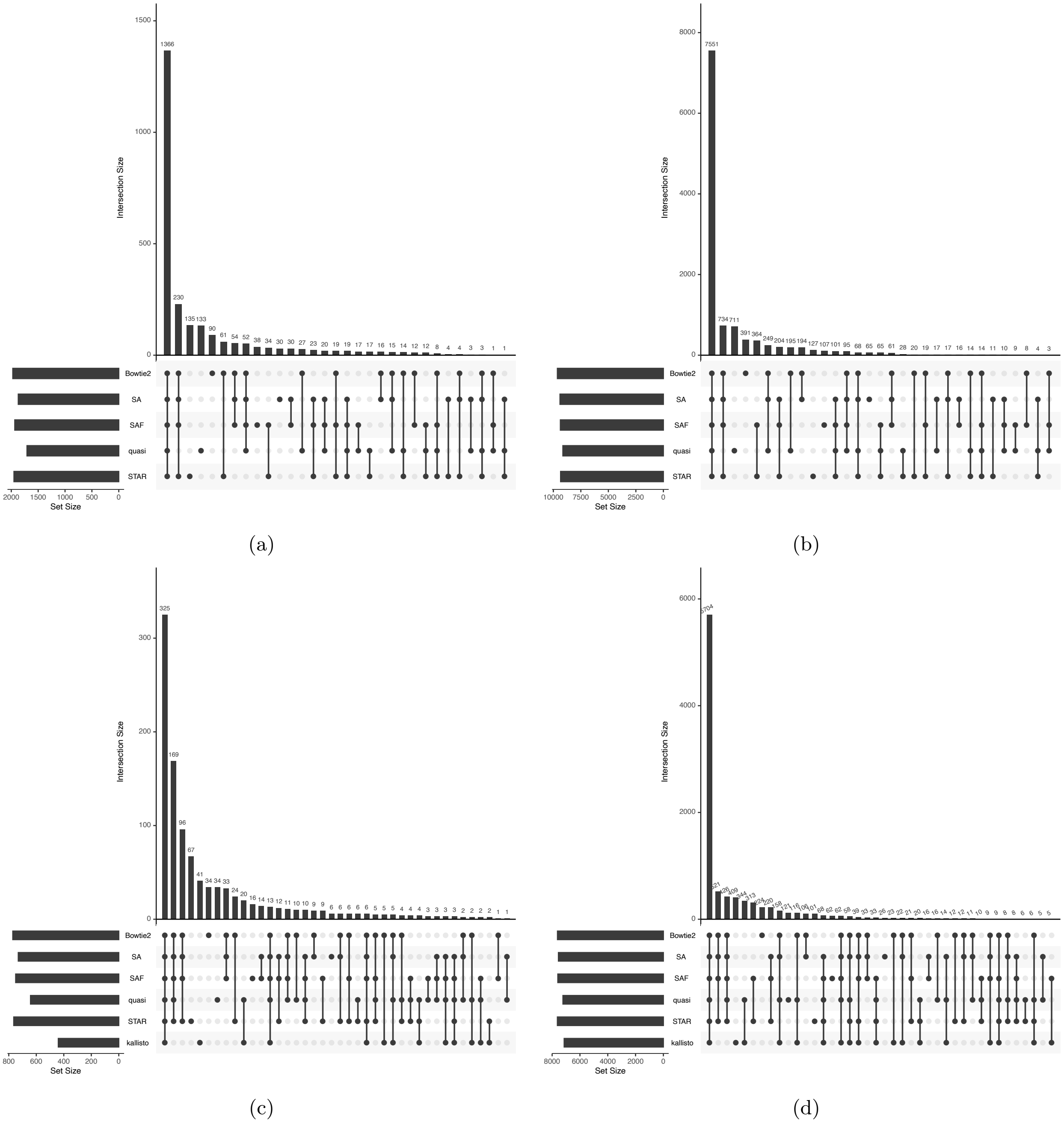


Figure S6: Comparison of sets of differentially expressed genes, and their overlaps, computed using each method. Figures (a) and (b) shows the results for the two datasets when filtered at an FDR of 0.05 and (c) and (d) shows the results at FDR 0.01 after including kallisto as an additional lightweight mapping approach.

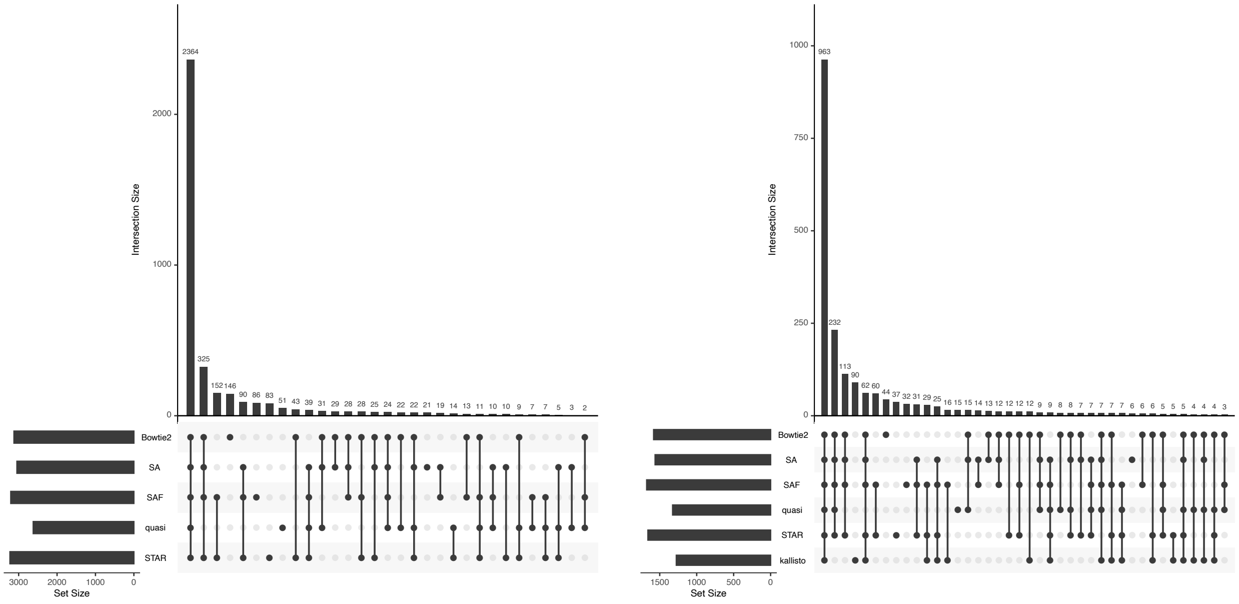


(a) (b)

Figure S7: Comparison of sets of differentially expressed transcripts, and their overlaps, computed using each method. Figure (a) shows the results when filtered at an FDR of 0.05 and (b) shows the results at FDR 0.01 after including kallisto as an additional lightweight mapping approach.
